# Attenuated estrogen signaling disrupts placentation and drives trophoblast defects in Down syndrome

**DOI:** 10.64898/2026.06.03.729739

**Authors:** Deirdre M. Logsdon, Isabela T. Pereira, Kindyll L. Wetta, Lauren Ohler, Mika Nevo, Bo D. Thorstenson, Brian F. Niemeyer, Bilge Birsoy, Lynae Smith, Cameron Hebert, John Rinn, Matthew D. Galbraith, Mary Allen, Robin Dowell, Joaquin M. Espinosa, Danny J. Schust, Justin Brumbaugh

## Abstract

Proper establishment of the primitive placenta and subsequent tissue homeostasis in the mature placenta are critical for successful pregnancy in humans. Placental insufficiency is associated with adverse pregnancy outcomes, including fetal growth restriction, preeclampsia, and pregnancy loss. Moreover, emerging evidence suggests that placental defects are associated with long-term health challenges that manifest well into adulthood; yet the etiologies of such diseases are largely unknown. Defining the mechanistic basis for placental deficiencies, therefore, has important implications for improving both reproductive health and the lifelong well-being of affected children. Down syndrome is characterized by placental defects of unknown mechanistic origin, and notably, individuals with Down syndrome are at increased risk of developing diseases commonly associated with placental insufficiency later in life. Using induced pluripotent stem cells from Down syndrome patients, we found that stem cell-based embryo models (i.e., blastoids) and directed differentiation systems recapitulate trophoblast cell fate defects observed in placentas affected by Down syndrome. Furthermore, we demonstrate that attenuated estrogen signaling contributes to placental syncytialization defects and identify NRIP1, a transcriptional corepressor of estrogen receptor that is located on chromosome 21, as a key driver of trophoblast cell fate defects. Increased gene dosage of NRIP1 in an otherwise diploid cell line phenocopies cell fate defects observed in trophoblasts affected by Down syndrome. Our study suggests that estrogen signaling is a crucial regulator of trophoblast development and may serve as a potential target for therapeutic intervention.

**Highlights and eTOC blurb:**

- Estrogen signaling mediates syncytiotrophoblast fusion
- Human iPS cells provide a tractable model for trophoblast defects in Down syndrome
- Trophoblast differentiation and estrogen signaling are disrupted in Down syndrome
- Increased NRIP1 expression is sufficient to induce trophoblast defects

Logsdon and colleagues apply patient-derived induced pluripotent stem cells to recapitulate placentation defects observed in Down syndrome. The authors demonstrate that attenuated estrogen signaling disrupts trophoblast differentiation and identify NRIP1, a gene found on chromosome 21 that dampens estrogen signaling, as a regulator of trophoblast maturation. NRIP1 and estrogen signaling may represent important therapeutic targets for infertility and Down syndrome.

## Introduction

The primitive placenta forms during embryo implantation to guide and support the developmental programming of a fetus through endocrine signaling, waste and nutrient exchange, and protection from insult. Inadequate establishment of the placenta during the first 12 weeks of gestation leads to clinically significant complications later in pregnancy, including fetal growth restriction, pre-term birth, preeclampsia, and pregnancy loss(1,2). Furthermore, a growing body of evidence suggests that inadequate placentation is a causal risk factor for the development of pathologies later in life, including obesity, diabetes, cardiovascular disease, and neurodegenerative disease(3,4). Despite the profound clinical impact of inadequate placentation on maternal and fetal health, the mechanisms that govern implantation and early placentation remain obscure.

Healthy human placentas comprise several trophoblast subtypes that arise from a progenitor population of cytotrophoblast (CTB) cells. CTB can either self-renew to replenish the stem cell pool or differentiate into one of two primary specialized cell types(5). Extravillous trophoblast (EVT) cells break away from fetal tissues and colonize the inner walls of the uterine blood vessels to ensure blood flow to the fetus. Meanwhile, syncytiotrophoblast (STB) cells serve as the primary contact with the maternal interface and mediate hormone production, as well as gas and nutrient exchange between mother and fetus. As part of their functional maturation, STB form from fused CTB to generate large, multinucleated syncytia, which secrete hCG and are readily detectable by microscopy. Together, these cell types maintain the crucial connection between a mother and child throughout pregnancy.

Although specialized trophoblast cells are critical in development and reproductive biology, substantial gaps remain in our understanding of the pathways that guide their differentiation and maturation. Factors that initiate EVT mobility and invasion are best characterized and include RHOA, ERK1/2, mTOR and others (reviewed in (6)). Functional maturation of STB cells likely involves canonical signaling pathways, including MAPK (7), Activin (8,9), and cyclic AMP/PKA (10,11), based on in vitro models derived from late-gestation and term placenta (reviewed in (11)). However, the signals that direct the formation of the primitive placenta and the origins of initiating signals are less clear. Specialized trophoblast cells are needed for implantation at approximately 6-9 days post-fertilization. This time point coincides with the elevation of tightly controlled endocrine signals from the ovary, such as estrogen, that prepare the uterus for implantation(12). Notably, estrogen receptor alpha is expressed in human trophoblast and increases during CTB differentiation (13) but is absent from mature STB late in gestation(14). The timing of these events raises the question of whether estrogen may also prepare trophoblast cells in the embryo for implantation. There is some evidence that estrogen acts directly on term placental explants (15); however, the mechanisms that control estrogen signaling in the primitive placenta are unknown.

STB defects severely compromise fetal development due to reduced nutrient availability, reduced secretion of placental hormones, and impaired invasion of the primitive trophoblast during the peri-implantation period (16). One example of such STB deficiency occurs in Down syndrome pregnancies. Down syndrome (trisomy 21, T21), resulting from a full or partial trisomy of the 21st chromosome, is the most common non-lethal aneuploidy disorder in humans and affects about 1 in 700 live births(17). Pregnancies affected by T21 are at increased risk for adverse pregnancy outcomes, including stillbirth and fetal growth restriction, both of which are associated with placental insufficiency(18). Individuals living with T21 are also at risk for various co-occurring conditions that critically affect their quality of life and lifespan, including obesity and congenital heart defects(19,20). Coincidentally, many co-occurring conditions observed in Down syndrome overlap with illnesses commonly associated with placental dysfunction. At present, STB defects have been observed in primary T21 cell cultures from mid- and late-gestation T21 placental tissue (21). But it is unclear if STB are affected in the primitive placenta, which forms in parallel with key organs and is therefore crucial for their development. Moreover, the underlying cause of STB defects in T21 remain unknown, in part due to the inaccessibility of early primary placental tissue and sample variation between human tissues, both of which limit precise functional studies or assays that require genetic manipulation.

To address these gaps, we applied trophoblast differentiation models using induced pluripotent stem (iPS) cell lines derived from an individual mosaic for trisomy of chromosome 21. We show that T21 trophoblast defects arise in a self-directed manner during implantation with stem cell-based embryo models. An orthogonal, directed differentiation system confirmed these defects and supported genomics studies, which revealed that T21 trophoblast-like cells have a dampened ability to respond to estradiol, highlighting an underappreciated role for estrogen signaling in the primitive placenta. Expanding our analysis to five additional pairs of iPS cell lines derived from donors with T21 and age-matched euploid controls, we confirmed these observations across various genetic backgrounds. Through further analysis, we identified nuclear receptor interacting protein 1 (NRIP1) as a contributing factor in defective STB cell fate commitment in T21 cells. Notably, NRIP1 is a co-transcriptional repressor of estrogen receptor and is located on chromosome 21, linking Down syndrome and defective regulation of estrogen signaling. Collectively, these studies point to NRIP1 and estrogen signaling as therapeutic targets for infertility, placental dysfunction, and Down syndrome.

## Results

### Down Syndrome Blastoids Display Implantation Defects

To model the formation of primitive placenta in the context of Down syndrome, we leveraged two iPS cell lines derived from a single chromosomally mosaic donor. One line is diploid for chromosome 21 (D21), and the other line is trisomic for chromosome 21 (Figure 1A, B), allowing us to directly compare T21 cells and a matching D21 control line that is otherwise isogenic. Using these lines, we developed stem cell-derived embryo models called blastoids that resemble pre-implantation blastocyst-like structures (Figure 1C). D21 and T21 cell lines formed blastoids with similar efficiency (Figure 1D) and stained positive for markers of embryonic and extra-embryonic lineages that are present in pre-implantation human embryos (Figure 1E and Figure S1A). We then transferred blastoids onto endometrial stromal cells to model implantation dynamics in vitro (22). After 3 days in extended culture on endometrial stromal cells, blastoids attached with clear trophoblast outgrowth (Figure 1F). Both T21 and D21 cultures were positive for human chorionic gonadotropin subunit beta (CGB), a hallmark of the multinucleated STB (Figure 1F and Figure S1B, C); however, CGB was decreased in the T21 line compared to D21 control (Figure 1G and Figure S1C). Moreover, we found that the number of nuclei within syncytia and the fusion index, a measure of functional maturation in STB, were significantly reduced in T21 outgrowths compared to D21 (Figure 1F, H), despite having similar outgrowth areas and total cell numbers (Figure S1D). Taken together, these data imply that fusion defects in T21 placentas may occur as early as the implantation stage.

**Figure 1.**
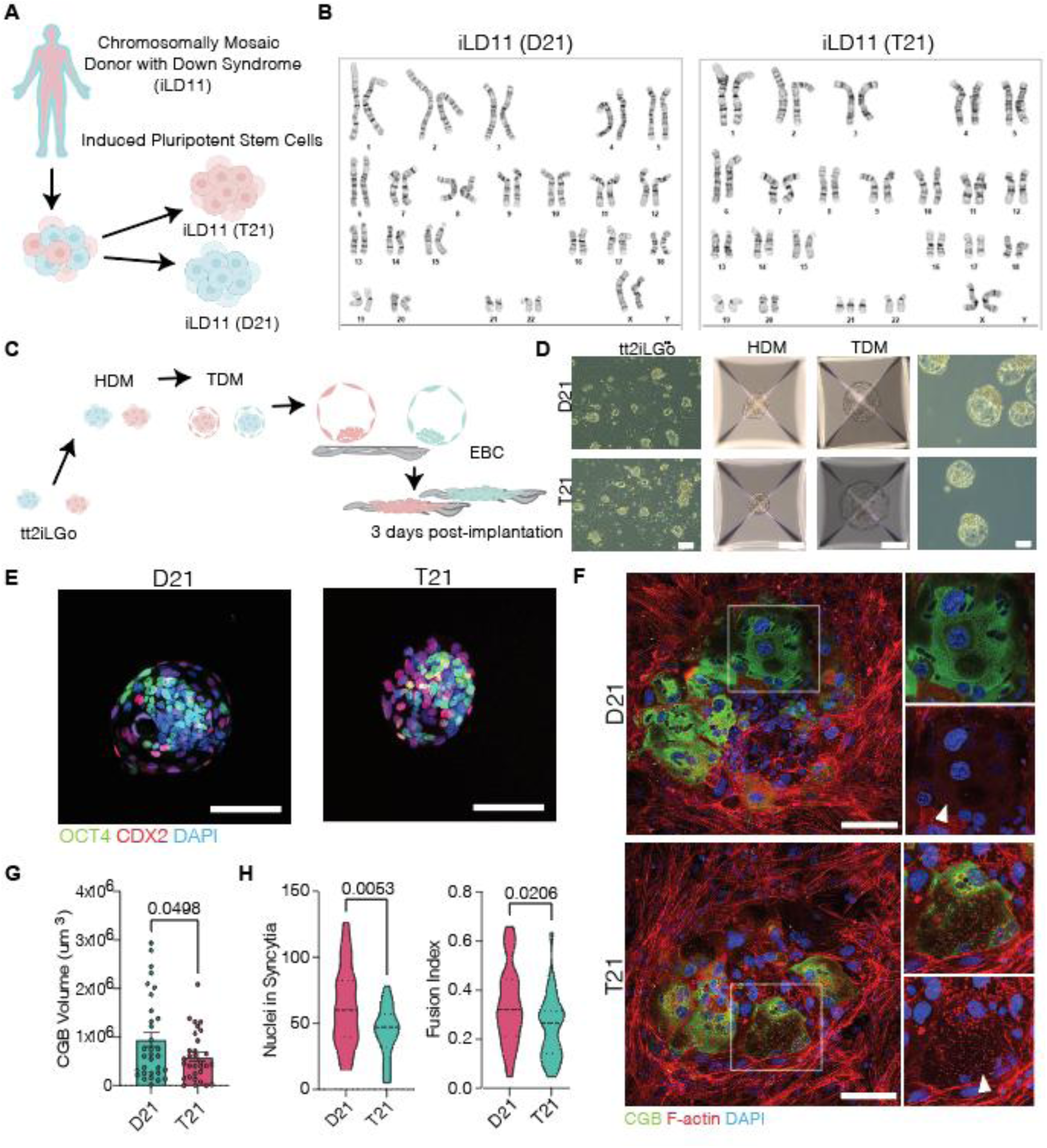
Down Syndrome Blastoids Display Differentiation Defects. A) Schematic of isogenic T21 and D21 iPS cell lines derived from a donor chromosomally mosaic for T21 (iLD11). B) Karyotype for isogenic T21 and D21 lines from mosaic donor iLD11. C) Schematic of blastoid generation from naïve pluripotent stem cells in tt2iLGo medium. HDM= hypoblast differentiation medium, TDM= trophoblast differentiation medium, EBC= extended blastoid medium. D) Representative brightfield images of T21/D21 isogenic iPS cells reset to a naïve pluripotent state and during blastoid generation. Scale bar=100um. E) Representative immunofluorescence images of T21/D21 blastoids stained for antibodies against pluripotency marker OCT4 (green), trophectoderm marker CDX2 (red), and DAPI (blue). Scale bar=100μm. F) Confocal microscopy of D21 (top) and T21 (bottom) blastoids 72h post implantation onto immortalized endometrial stromal cells. CGB=green, F-actin=red, DAPI=blue. Scale bar=100um. G) Volume of human chorionic gonadotropin beta (CGB) signal after immunofluorescence in T21/D21 implanted blastoids. H) Number of nuclei in syncytia (left) and fusion index (right, (number of nuclei in syncytia-number of syncytia)/total nuclei). P values are represented for each comparison following student t test.

### Directed Differentiation Models Define STB Differentiation Defects in T21 Cells

To scale our approach for detailed molecular analyses, we converted the matching iPS cell lines into a trophoblast-like fate by applying BMP4, A83-01 (a TGFβ inhibitor), and PD173074 (an FGFR2 inhibitor) for 8 days(23). Following differentiation, iPS cells gave rise to a mixed population of mono-nucleated CTB-like and multinucleated STB-like cells (Figure 2A) that stained positive for pan-trophoblast markers cytokeratin-7 (*KRT7*), GATA binding protein 3 (*GATA3*) (Figure S2A), and secrete reduced human chorionic gonadotropin (hCG) into spent medium (Figure 2B). Notably trophoblast-like cells derived from this process have a gene expression profile that is distinct from late-stage placenta but largely overlaps with first-trimester trophoblasts and peri-implantation stage trophoblasts from implanting embryos (24). Thus, STB that emerge from this system likely represent the primitive syncytium. Comparing T21 to D21 samples, we observed decreased nuclei in syncytia and decreased fusion indices in T21 samples (Figure 2C, D), as well as decreased hCG production (Figure 2B), confirming STB differentiation defects in Down syndrome samples in an orthogonal differentiation system.

**Figure 2.**
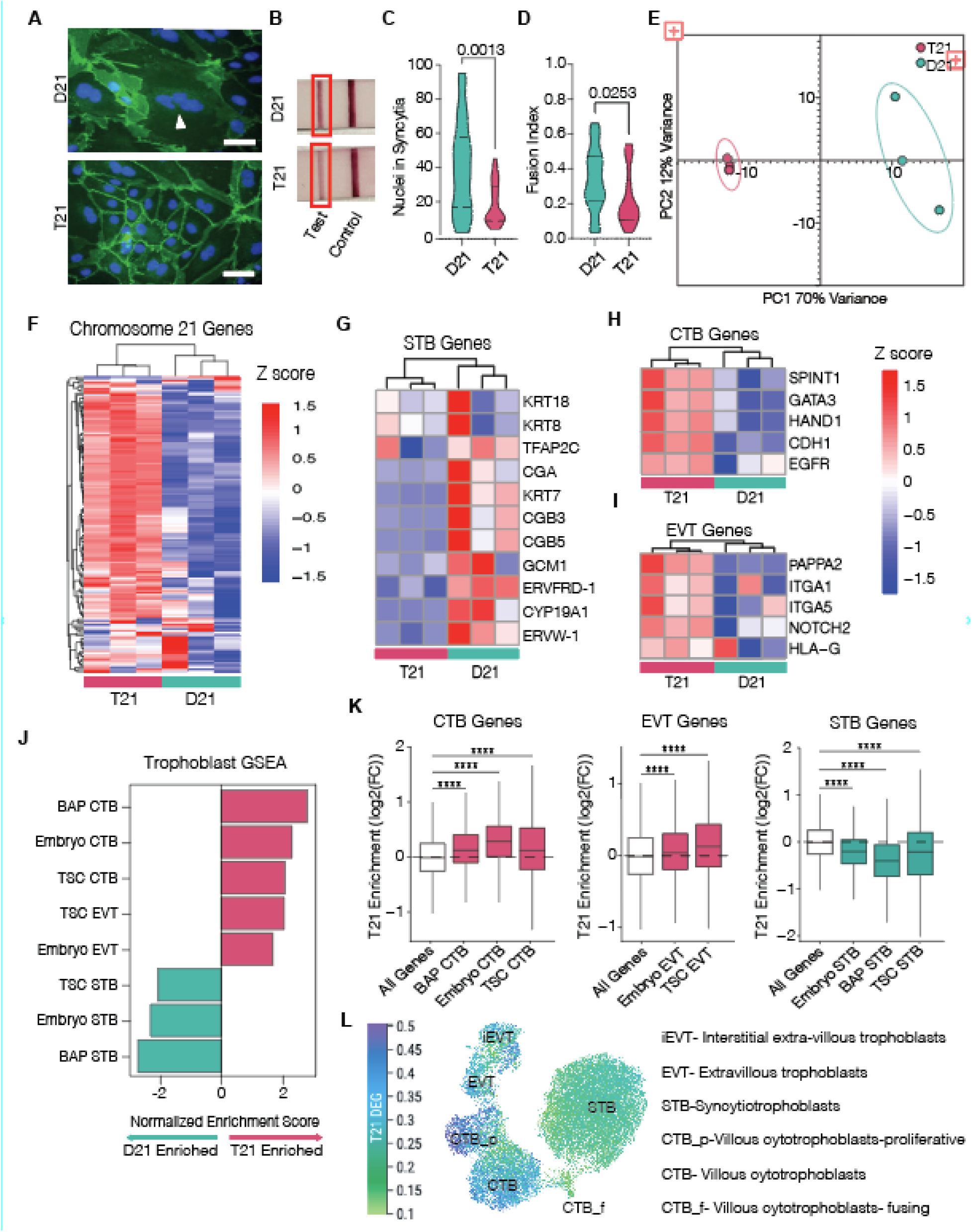
Down Syndrome Trophoblast-like Cells Phenocopy Fusion Defects. A) Representative immunofluorescence microscopy for D21 or T21 trophoblast-like cells following differentiation. DAPI=blue, E-cadherin=green. White arrows highlight multinucleated cells with 2 or more nuclei. Scale= 100µm. B) Pregnancy test strips for secreted human chorionic gonadotropin from spent medium for D21/T21 trophoblast-like cells. C) Nuclei in syncytia and D) fusion index as quantified by (nuclei in syncytia-number of syncytia)/total nuclei per field of view *p<0.05, **p<0.01. n=10 fields of view for each of 3 individual replicates E) PCA plot. n=3 individual replicates. F) Heatmap with Z-score scaled expression of genes on human chromosome 21 in D21/T21 isogenic trophoblast-like cells. G) Heatmap with Z-score scaled expression of STB genes, H) CTB, and I) EVT genes in D21/T21 isogenic trophoblast-like cells.). J) GSEA for CTB, STB, and EVT gene sets from peri-implantation embryos(55), BMP4-treated trophoblast-like cells (BAP), and trophoblast stem cells (TSC)(25) with normalized enrichment score in T21/D21 trophoblast-like isogenic samples. K) Boxplots showing log2 fold change for trophoblast gene sets from panel I. ****p<0.0001 following Welch’s t. test comparing each gene set to all other genes. L) Uniform manifold approximation and projection (UMAP) plot of placental implantation site of a donor from previously published single nuclei RNA-seq with 37,675 nuclei (5) showing expression of T21 DEG.See also Figure S1, S2.

To characterize gene expression in T21 and D21 trophoblast cultures, we performed RNA-sequencing (RNA-seq) analysis on our isogenic lines after 8 days of differentiation in BMP4, A83-01, and PD173074. As expected, T21 samples clustered separately from D21 by principal component analysis (PCA) (Figure 2E). These differences reflect, in part, increased dosage of genes on chromosome 21 (Figure 2F); however, we noticed that many genes associated with STB (e.g., *CGB3/5, CGA, GCM1, ERVW-1)* were depleted from T21 trophoblast-like cells (Figure 2G). By contrast, CTB genes (e.g., *GATA3, CDH1, HAND1*) and EVT genes (e.g., *HLA-G, ITGA5, NOTCH2*) were increased in T21 cells relative to D21 cells (Figure 2H, I). To extend this analysis using a more comprehensive, unbiased approach, we performed gene set enrichment analysis (GSEA)(25) and confirmed that transcriptional signatures associated with STB were significantly depleted in T21 lines, whereas CTB and EVT signatures were enriched in T21 lines (Figure 2J, K). These data are consistent with defects observed in culture and suggest a differentiation bias in T21 cells. To further refine this bias, we generated a list of differentially expressed genes (DEGs) between T21 and D21 trophoblast-like cells. We then examined their expression patterns in trophoblast subtypes, based on published single nuclei RNA-seq data from placental tissue at 8-9 weeks gestation(5). Notably, STB and CTB_F, which represents a fusing STB-differentiation intermediate, were enriched for genes downregulated in T21 cells compared to D21 controls (Figure 2L). These data suggest that genes related to STB fate fail to activate early in the differentiation process of T21 trophoblast-like cells. Genes upregulated in T21 cells were enriched to varying degrees in EVT and CTB, but were particularly high in CTB_P, a proliferating population of trophoblast progenitor cells. This was also true for gene sets generated from single cell RNA-seq data from placentas from 6-14 gestational weeks and single cell RNA-seq data from chorionic villi at 6-11 weeks of gestation (5,26,27)(Figure S2B). Together, these data suggest that trisomy 21 leads to impaired trophoblast differentiation that favors CTB self-renewal and, to a lesser extent, EVT differentiation at the expense of STB.

EVT cells are highly mobile, which provides a tractable way to test their functional maturation in vitro. We therefore set out to quantify EVT mobility in T21 and D21 cells; however, our prior two-dimensional differentiation system generates a mixed population of trophoblast cells (28), making direct functional assays challenging. To address this issue, we adapted a BMP4/Sphingosine 1-phosphate-driven differentiation protocol that generates a higher proportion of EVT(29) and confirmed the expression of EVT marker *MMP2* (Figure S2C). We then performed scratch wound assays to test EVT mobility. In line with our gene expression analysis, T21 EVT-like cells achieved nearly 60% wound closure by 6 hours, while D21 EVT-like cells reached only ∼30% wound closure at the same time point (Figure S2, E). These results suggest that T21 EVT are functionally mature and highly mobile compared to D21 controls and further support the notion that trisomy 21 biases cell fate in trophoblast.

### Trisomy 21 STB Defects are Consistent Across Genetic Backgrounds, but Severity Varies

Individuals with Down syndrome display a continuum of disease severity, potentially due to genetic background and variegated expression of genes on chromosome 21(24). To test whether trophoblast defects were consistent across different genetic backgrounds, we expanded our analysis to include five additional iPS cell lines derived from T21 donors, each with an age- and sex-matched D21 control (Figure S2F). These lines span age groups from 6 years old to 47 years old at the time of collection and include cells derived from both male and female donors. We then induced differentiation of each line into trophoblast-like cells and quantified fusion indices as a proxy for STB differentiation efficiency. Consistent with analysis from isogenic lines, fusion indices were significantly decreased in four of five T21 lines (Figure S2G-K). The final cell line also showed a decrease in fusion index but was more variable and did not reach statistical significance (Figure S2G). Thus, STB defects in T21 trophoblast-like cells were generally consistent across genetic backgrounds.

To extend our findings, we performed RNA-seq in trophoblast-like cells and undifferentiated iPS cells for all 6 pairs of age- and sex-matched T21/D21 cell lines, including the isogenic lines. As expected, undifferentiated iPS cells expressed pluripotency transcripts, including *SOX2*, *NANOG*, and *POU5F1,* at high levels compared to trophoblast-like cells (Figure S3A). Likewise, trophoblast-like cells expressed genes associated with trophoblast subtypes, as well as pan trophoblast markers (Figure S3A). Considering all 6 pairs of lines, a few individual genes were consistently differentially expressed (Figure S3B), likely due to variability between genetic backgrounds. This variability was reflected by PCA, which revealed no clear clustering of D21 and T21 lines in either iPS or trophoblast-like cells (Figure S3C), unless the PCA was limited to genes on chromosome 21 (Figure S3D). Despite this variability, GSEA revealed that CTB and EVT transcriptional signatures were again enriched in T21 lines, while STB gene expression profiles were largely reduced in T21 lines (Figure S3E), in line with decreased fusion indices detected in culture. Moreover, STB-associated genes were largely decreased in T21 lines compared to D21 controls (Figure S3F). Together, these data suggest that although few genes are significantly differentially expressed across genetic backgrounds, overall transcriptional signatures important for STB maturation are consistently disrupted in T21 trophoblast-like cells.

### T21 Trophoblast-like Cells Have Impaired Estrogen Signaling

We next sought to identify potential pathways that are altered in T21 cells and may interfere with STB differentiation. To eliminate variability from genetic background, we considered gene expression from the isogenic lines. In total, T21 trophoblast-like cells had 2476 DEGs compared to D21 (Figure 3A). To identify functional categories altered in T21 trophoblast-like cells compared to D21 control, we performed GSEA using a curated set of hallmark gene sets. T21 trophoblast-like cells showed enrichment of the interferon (IFN) alpha and gamma responses, in alignment with previous reports showing that T21 consistently activates the IFN response across multiple cell types, including embryonic tissues(30–32). By contrast, transcriptional profiles depleted from T21 trophoblast-like cells included estrogen-response signatures, suggesting that estrogen signaling may be dampened in T21 trophoblasts (Figure 3B).

**Figure 3.**
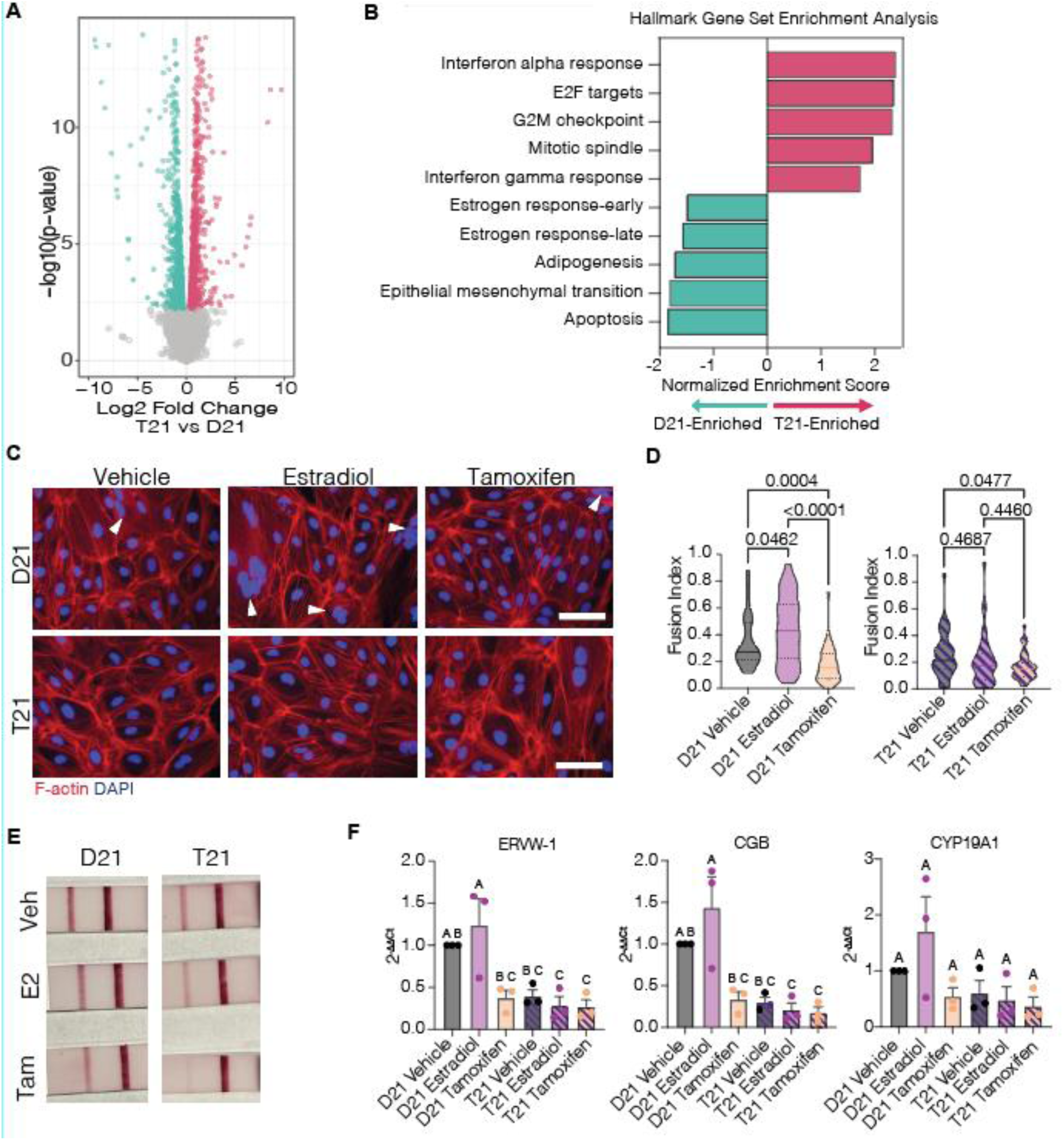
Down Syndrome Trophoblast-like Cells Have Attenuated Estrogen Signaling. A) Volcano plot highlighting differentially expressed genes between T21 (red) or D21 (left) isogenic trophoblast-like cells. RNA-seq from 3 individual replicates of isogenic iLD11 iPS cells. B)GSEA highlighting hallmark gene sets significantly enriched in T21 or D21 isogenic trophoblast-like cells. C) Representative immunofluorescence microscopy for D21/T21 trophoblast-like cells following differentiation in either vehicle, 1µM 17β-estradiol, or 1µM Tamoxifen. DAPI=blue, F-actin=red. White arrows highlight multinucleated cells with 3 or more nuclei. Scale= 100µm. D) Fusion indices for D21 or T21 trophoblast-like cells when treated with vehicle, 1µM 17β-estradiol, or 1µM Tamoxifen. Significance is noted following a one-way ANOVA with Tukey’s multiple comparisons test. n= 40 fields of view per treatment across 4 individual replicates. E) Pregnancy test strips for secreted human chorionic gonadotropin from spent medium for D21/T21 trophoblast-like cells treated with 17β-estradiol (E2) or Tamoxifen (Tam). F) Relative transcript expression following quantitative RT-PCR for STB transcripts *ERVW-1*, *CGB*, and *CYP19A1*. Significance is noted p<0.05 with different letters following a one-way ANOVA with Tukey’s multiple comparisons test. Data are represented as mean ± SEM. See also Figure S3.

Previous work in trophoblast explants derived from term placenta suggested that stimulating estrogen increases syncytialization(15). Moreover, genes important for STB fusion, including *ERVW-1*, contain estrogen response elements in their upstream cis-regulatory sequences, providing a potential mechanistic link between estrogen signaling and STB formation. To test this possibility, we treated isogenic T21 or D21 cells with 1µM 17 β−estradiol to activate the estrogen receptor, or 1µM Tamoxifen, an estrogen receptor antagonist, and monitored fusion index as a measure of STB differentiation. D21 cells increased cell fusion when treated with 17 β−estradiol, while Tamoxifen significantly disrupted the formation of multinucleated STB to a level commensurate with T21 cells (Figure 3C, D). By contrast, T21 trophoblast-like cells were refractory to 17 β−estradiol stimulation, suggesting that estrogen signaling is attenuated in Down syndrome. Correspondingly, the expression of genes related to STB and estrogen response increased in D21 cells following 17 β−estradiol treatment and decreased with Tamoxifen to a level similar to T21 cells, which were unaffected by either treatment. This pattern was likewise reflected in hCG secretion, suggesting that estrogen signaling is suppressed in T21 cells (Figure 3E, F).

To identify potentially causal genes on chromosome 21 that may influence estrogen responsiveness, we filtered DEGs between T21 and D21 trophoblast-like cells for transcriptional regulators. Of the approximately 233 protein-coding genes on chromosome 21, 68 were significantly differentially expressed between Down syndrome and diploid lines, and of these genes, 18 are classified as transcriptional regulators (Figure 4A, B). Primary among them is Nuclear Receptor Interacting Protein 1 (*NRIP1*) (Figure 4C), which is a known transcriptional co-repressor for estrogen receptor (33). Given that estrogen signaling is attenuated in T21 trophoblast-like cells, we hypothesized that increased gene dosage of *NRIP1* may disrupt STB differentiation, dampening estrogen-regulated transcription.

**Figure 4.**
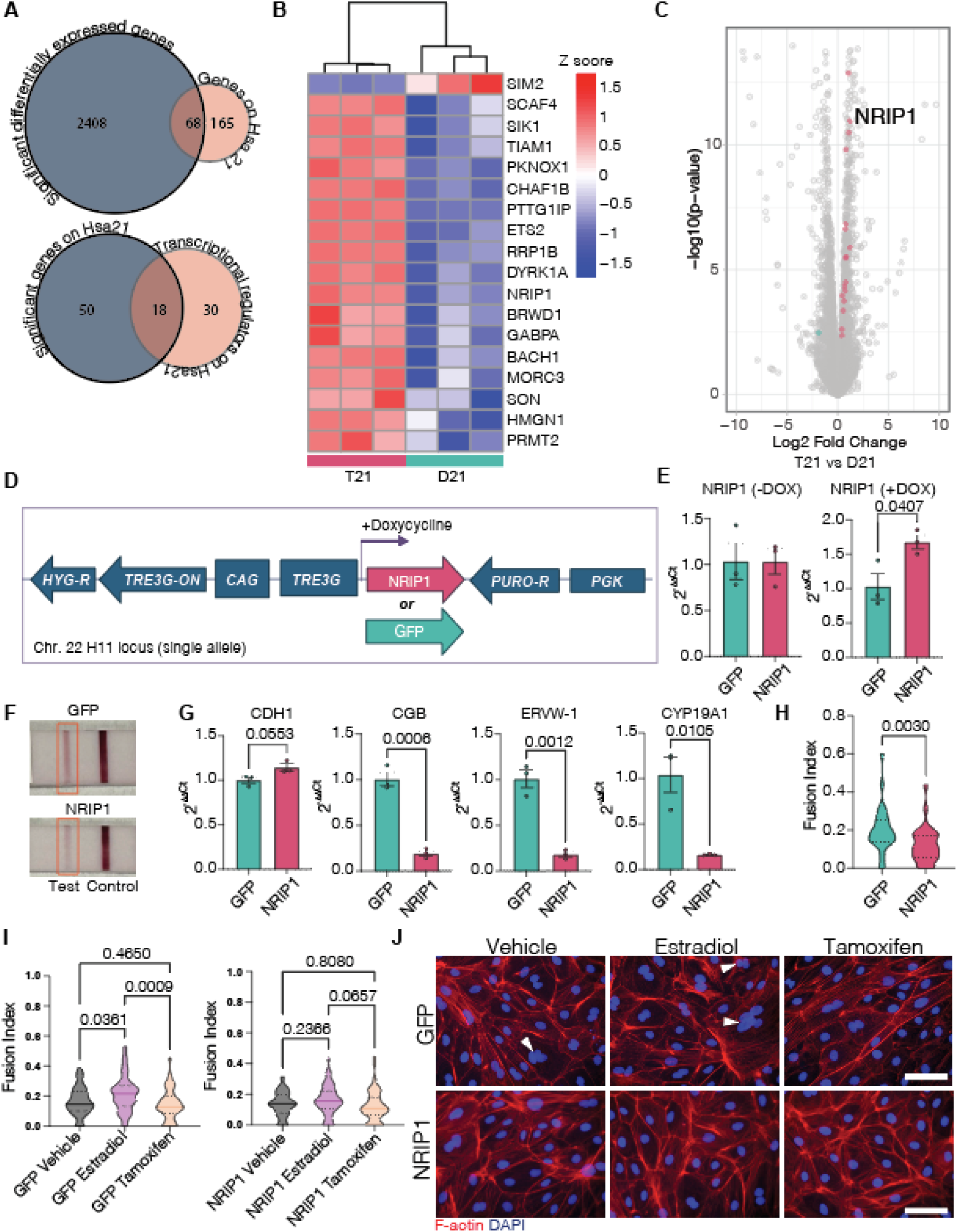
Overexpression of NRIP1 Disrupts Differentiation of Trophoblast-like Cells. A) Venn diagrams highlighting 68 significantly differentially expressed protein coding genes on chromosome 21 (top) and significant transcriptional regulators on chromosome 21 (bottom). B) Heatmap with Z-score scaled expression of transcriptional regulators on chromosome 21 differentially expressed between D21/T21 isogenic trophoblast-like cells. C) Volcano plot highlighting differentially expressed transcriptional regulators on chromosome 21 between T21 (red) or D21 (left) isogenic trophoblast-like cells. D) Schematic of NRIP1 or GFP insertion into the H11 safe harbor locus on chromosome 22 under control of doxycycline. E) NRIP1 transcript expression without doxycycline ( -DOX, left) or in 2ug/mL doxycycline (+DOX, right). F) Pregnancy test strips for secreted human chorionic gonadotropin from spent medium for GFP/NRIP1 trophoblast-like cells. G) Relative transcript expression following quantitative RT-PCR for STB transcripts *ERVW-1*, *CGB*, and *CYP19A1* and cytotrophoblast transcript *CDH1*. *p<0.05, **p<0.01, ***p<0.001 following student T. test. Data are represented as mean ± SEM. F) Fusion index as quantified by (nuclei in syncytia-number of syncytia)/total nuclei per field of view **p<0.01 following student T. test. n= 30 fields of view across 3 individual replicates. I) Fusion indices for GFP or NRIP1 overexpression trophoblast-like cells when treated with vehicle, 1µM 17β-estradiol, or 1µM Tamoxifen. Significance is noted following a one-way ANOVA with Tukey’s multiple comparisons test. n= 60 fields of view across 3 individual replicates. J) Representative immunofluorescence microscopy for D21/T21 trophoblast-like cells following differentiation in either vehicle, 1µM 17β-estradiol, or 1µM Tamoxifen. DAPI=blue, rhodamine phalloidin=red. White arrows highlight multinucleated cells with 2 or more nuclei. Scale= 100um. See also Figure S4.

To test this possibility, we modified a diploid iPS cell line by integrating an additional copy of *NRIP1* under the control of a doxycycline-inducible promoter into the H11 safe-harbor locus on chromosome 22 (Figure 4D). In parallel, we generated a matching GFP control line (Figure 4D). Doxycycline induction elevated *NRIP1* ∼1.5-fold in the *NRIP1* overexpression line, matching expression levels expected in T21 cells (Figure 4E). Next, we initiated differentiation into trophoblast-like cells and measured fusion indices to determine differentiation efficiency. Notably, increased *NRIP1* expression suppressed cell fusion, similar to T21 lines (Figure 4F). Expression of estrogen-responsive and STB-related transcripts were similarly decreased in the *NRIP1* overexpression line, as was hCG secretion (Figure 4G, H). Finally, *NRIP1* overexpressing cells failed to respond to estradiol treatment, similar to T21 cell lines (Figure 4I, J). Collectively, these data suggest that increasing *NRIP1* transcript levels 1.5-fold is sufficient to dampen estrogen signaling in trophoblasts and disrupt STB fusion.

## Discussion

The placenta is fundamentally important for human pregnancy; however, the mechanisms that govern early trophoblast development during implantation remain elusive. Understanding these mechanisms is crucial for treating pregnancy complications and diseases associated with placental insufficiency, including Down syndrome. Despite its importance to both maternal and fetal health, studying the primitive placenta is challenging. Primary human tissue representing the primitive placenta is difficult or impossible to obtain, given both ethical and practical constraints. Animal models are an option, but placental physiology differs dramatically between species, limiting the relevance of studies in common animal systems such as mice(34). Moreover, in the context of Down syndrome, genomic organization differs between mouse and human such that genes present on human chromosome 21 are scattered across multiple chromosomes in mouse. As a result, mouse models of T21 contain a majority, but not all orthologs for human T21 genes(35). For example, *NRIP1*, identified in this study as a contributor to placental defects in T21 lines, is not trisomic in the commonly used Ts65Dn mouse. Therefore, careful mechanistic studies on T21 placentation must be performed in physiologically relevant human trophoblast systems.

Here, we leveraged age- and sex-matched D21 and T21 iPS cells as experimental systems to study early trophoblast development and to assess STB fusion and differentiation defects in Down syndrome placentation. Multiple, orthogonal approaches confirmed that T21 iPS cells recapitulate placental defects observed in vivo. Notably, blastoids represent a self-organizing system that permits investigation of implantation and primitive placenta formation, events that are otherwise difficult to capture experimentally. To our knowledge, this is the first study to use blastoids to model a disease phenotype associated with a complex aneuploidy.

A fundamental challenge in studies with multiple human cell lines is transcriptional and phenotypic variability inherent to different genetic backgrounds. Indeed, we identified few common differentially expressed genes when analyzing all six pairs of age- and sex-matched lines. Despite this variability, functional and gene expression defects in STB differentiation emerged as common between T21 cell lines. By then focusing on T21 and D21 iPS cell lines derived from a single, chromosomally mosaic individual, we eliminated genetic background as a variable between lines. Our analysis revealed that attenuated estrogen signaling through increased gene dosage of *NRIP1* contributes to placental dysfunction in Down syndrome. NRIP1 functions as a negative coregulator of estrogen receptor signaling(36–40). Estrogen secretion by the ovary increases during the growth of a pre-ovulatory follicle, preceding a sharp drop after ovulation. Following this decrease, there is a secondary, less pronounced “nidatory” wave of estrogen alongside a rise in progesterone during the window of implantation. Together, progesterone and estrogen act on the endometrium to prepare the uterus for implantation, and deviations in this ratio can be deleterious (12)(41). However, the direct role of estrogen on the embryo is less clear. Estradiol levels above 100nM negatively impact embryo implantation in mice (42), but mouse blastocysts treated with an estrogen receptor inhibitor also have reduced pre- and peri-implantation dynamics in vitro (43). This implies that embryos operate in a “Goldilocks zone” in which the nidatory wave of estrogen must fall within an optimal level. To maintain a precisely-controlled estrogen response, both positive and negative feedback mechanisms likely exist. We propose that NRIP1 operates as a negative regulator of estrogen signaling in the primitive placenta. This is particularly informative for human assisted reproductive technology applications, as estrogen supplementation into pre-implantation blastocyst medium is largely unexplored, and exogenous estrogen supplementation during the window of implantation is a topic of debate with conflicting data (12,44). Our work highlights the importance of estrogen signaling directly on STB fusion and proposes a novel role of NRIP1 in maintaining a tightly controlled estrogen response in the primitive placenta.

While our work suggests an important role for estrogen signaling and *NRIP1* in STB differentiation, other genes are required for cell fusion, including *ERVW-1* and *ERVFRD-1* (45,46). Moreover, several genes on chromosome 21, including *ERVH48-1,* likely play a role in defective STB fusion in T21 (47–49). Understanding the precise role of these factors and their interplay in the context of placentation in Down syndrome will be important moving forward.

### Experimental procedures

#### Ethics Statement

Research involving the use of induced pluripotent stem cell lines for the production of blastoids and subsequent implantation was approved by an institutional Scientific Ethics Committee for the University of Colorado at Boulder through the Environmental Health and Safety Office. All blastoid experiments were completed in compliance with the 2021 International Society for Stem Cell Research Guidelines for Stem Cell Research and Clinical Translation (50). All blastoid experiments were stopped after 3 days post implantation which is equivalent to approximately 8 d. p. f. in human embryos well within the 14-day rule. Induced pluripotent stem cell lines were obtained from the Human Trisome Project biobank administered by the Linda Crnic Institute for Down Syndrome under an IRB protocol overseen by the Colorado Multiple Institutional Review Board (COMIRB #15-2170). Donors or their legal guardians provided consent for biospecimen collection and iPS cell generation. All blastoid reagents and experiments were supported by non-federal funding through the Anna and John J. Sie Foundation and institutional startup funds from the University of Colorado. All other experiments were funded through the National Institutes of Health NRSA F32 award number 1F32HD116568-01 and NIH R21 award number 1R21HD118444-01.

#### Statistics

All statistical analyses were generated with GraphPad Prism version 11.0.0. and R version 4.5.3. All experiments with continuous datapoints are represented as mean ± SEM and were evaluated for statistical significance following a student t test for two comparisons, or a one-way ANOVA with Tukey’s multiple comparison test with three or more groups. Significance was defined as p<0.05. Normality was assessed with a Shapiro-Wilk test for normality. If assumptions of normality were not met, a Mann-Whitney or Kruskal-Wallis test for nonparametric comparisons was used. For RNA-seq experiments, donor pairs were treated as a paired variable wherever possible, and statistical comparisons were performed within matched donor pairs to reduce inter-individual variability.

#### Induced Pluripotent Stem Cell Lines

iPS cell lines were obtained from the Human Trisome Project Biobank (www.trisome.org) administered by the Linda Crnic Institute for Down Syndrome at the University of Colorado Anschutz. iPS cell lines were reprogrammed from either peripheral blood monocytes with a non-integrating episome (iLC68BC-5 and iLD49BC-3) or from renal epithelial cells with a 5fM3O mRNA cocktail (iLC42-3, iLC47-4, iLC62-6, iLC67-3, iLD11, iLD19-4, iLD2-3, and iLD23-2). Primed iPS cell lines were stably maintained on Matrigel (Corning) in essential 8 (E8) medium (DMEM/F12, 20 µg/mL insulin, 10.7 µg/mL transferrin, 100 ng/mL human basic fibroblast growth factor 2 (FGF2), 2 ng/mL transforming growth factor beta (TGFβ), 14 ng/mL sodium selenite, 64 µg/mL L-ascorbic acid-2-phopshate). Medium was exchanged daily and cells were passaged with 0.5mM ethylenediaminetetraacetic acid (EDTA) approximately every 3-4 days. Trisomy 21 was verified by karyotype (Figure 1B) after initial derivation and were regularly checked for maintenance of trisomy 21 via quantitative RT-PCR of genomic DNA with primer pairs for chromosome 1 and chromosome 21 (Table 1).

**Table 1.**
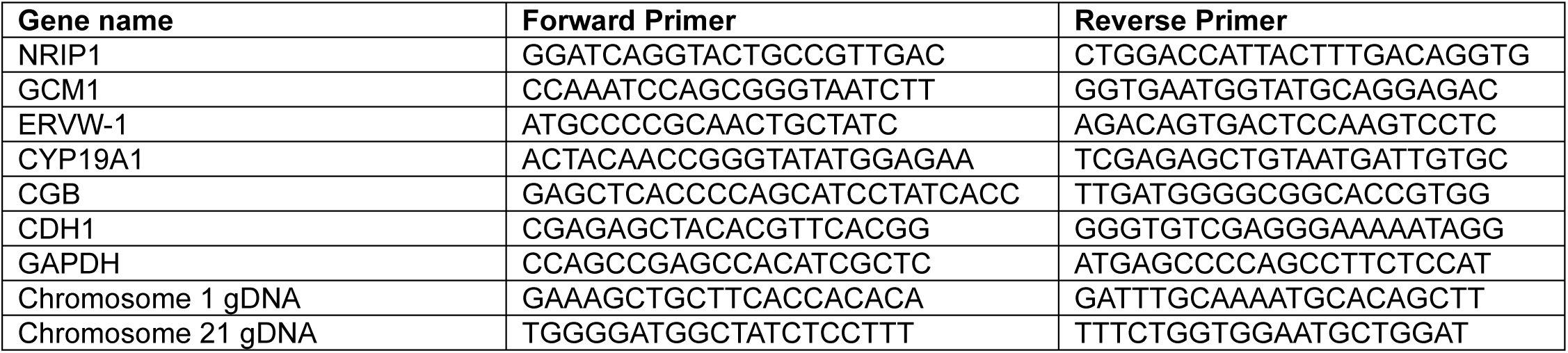
Primers.

#### Generation of NRIP1/GFP Overexpression Lines

NRIP1 coding sequence was synthesized by Twist Bioscience and cloned into the linearized pCR-XL-lox511-PURO-lox2271 vector using In-Fusion Cloning (Takara Bio) according to the manufacturer’s instructions. The construct was verified by sequencing. Trisomic NRIP1 overexpression line was generated from a human-induced pluripotent stem cell line (Applied Stem Cell, ASE-9211) engineered to include a dox-inducible expression cassette and Lox sites within the H11 locus on one single allele. The NRIP1 or GFP coding sequence was inserted into the dox-inducible expression cassette alongside a puromycin resistance gene. For transgene integration, iPSCs were dissociated using Accutase and seeded onto Geltrex-coated 6-well plates at a density of 3.5 × 10^5^ cells per well in medium supplemented with CEPT Cocktail (MedChemExpress, HY-K1043). After 2 hours, cells were transfected with 300 ng of Cre plasmid and 1800 ng of NRIP1 construct plasmid using 7.5 μL of Lipofectamine Stem Transfection Reagent (Invitrogen, STEM00008). When the transfected cultures reached 70-80% confluency, cells were split into Geltrex-coated 12-well plates supplemented with 10 μM Y-27632. Puromycin selection started on the following day with 200 ng/mL Puromycin (InvivoGen, ant-pr-1), and selection medium was refreshed daily until distinct single colonies appeared. Individual colonies were identified and manually picked under a microscope. Stable cell lines were derived from each colony, expanded, and cryopreserved. NRIP1 overexpression was validated via RT-qPCR after exposure to 100ng/mL doxycycline (Sigma, D5207) for 24h.

#### Naïve Resetting, Blastoid Generation, and Extended Culture of Blastoids

Isogenic iPS cell lines were reset to a naïve pluripotent state on mitotically inactive mouse embryonic fibroblasts treated with mitomycin C(51). Cells were first treated sequentially with resetting medium 1 (DMEM/F12, 0.5x N2, 0.5x B27, 1x GlutaMAX, 0.1mM 2-mercaptoethanol, 1 mM valproic acid, 1 µM PD0325901, 10 ng/mL human leukemia inhibitory factor (hLIF)) for 3 days, resetting medium 2 (DMEM/F12, 0.5x N2, 0.5x B27, 1x GlutaMAX, 0.1mM 2-mercaptoethanol, 1 µM PD0325901, 10 ng/mL hLIF, 2 µM XAV939, 2 µM Go6983) for 5 days, and maintained for up to 6 passages in tt2iLGo (DMEM/F12, 0.5x N2, 0.5x B27, 1x GlutaMAX, 0.1mM 2-mercaptoethanol, 1 µM PD0325901, 10 ng/mL hLIF, 2 µM XAV939, 2 µM Gö6983, 1µM CHIR99021). Only naïve cells with a domed morphology and between 3-7 passages in tt2iLGö were used for blastoid generation.

To generate pre- and peri- implantation stage blastoids, we used an established protocol (22). In brief, naïve cells were seeded into Aggrewell dishes at ∼25,000 cells per microwell. Cells were initially seeded in hypoblast differentiation media (50% DMEM/F12, 50% Neurobasal, 1x GlutaMAX, 0.1mM 2-mercaptoethanol, 1x NEAA 1x N2, 1x B27, 20ng/mL FGF2, 20 ng/mL Activin A, 3µM CHIR99021, 50nM Chroman1, 5 µM Emricasan, 1x Polyamines, 0.7µM Trans-ISRIB) and centrifuged at 500 rcf for 3 minutes. Each day half of the medium volume was exchanged with fresh medium. On day 3 after initial seeding, full medium was exchanged for trophoblast differentiation medium (75% DMEM/F12, 25% Neurobasal, 0.5x GlutaMAX, 0.25x N2, 0.25x B27, 0.5x NEAA, 0.1mM 2-mercaptoethanol, 1µM PD0325901, 1µM A83-01, 0.5µM SB590885, 1µM WH-4-023, 10ng/mL hLIF, 0.5µM LPA, 0.5x ITS-X, 50nM Chroman1, 5µM Emricasan, 1x Polyamines, 0.7µM Trans-ISRIB). Half volumes were exchanged daily for 5 additional days, and blastoids were harvested at day 8. Blastoids selected for downstream analysis were chosen if <30% of their total volume was ICM-like with ∼60% blastocoele-like morphology and ∼150µm in diameter.

Selected blastoids were seeded 3 per well onto mitotically inactive immortalized endometrial stromal cells from a patient biopsy in extended blastoid medium (50% DMEM/F12, 50% Neurobasal, 1x GlutaMAX, 0.1mM 2-mercaptoethanol, 1x NEAA 1x N2, 1x B27, 8nM estradiol, 200ng/mL progesterone, 1mM sodium pyruvate, 50nM Chroman1, 5µM Emricasan, 1x Polyamines, 0.7µM Trans-ISRIB) pre-plated onto chambered coverslips (ibidi). Blastoids were left undisturbed for 3 days. Peri-implantation blastoid culture experiments were performed with 9-10 blastoids over 3 individual replicates.

#### Differentiation of Trophoblast-like Cells

Cells were passaged at least once following thaw and were differentiated with an established protocol with minor modifications (23). Primed-state iPS cells were seeded in E8 medium on Matrigel following EDTA dissociation in small clumps (∼5-10 cells) at about 100,000 cells/9.5cm^2^. The following day (day -1), medium was exchanged for a simple transition medium eliminating TGFβ (DMEM/F12, 4ng/mL FGF2, 20% Knockout Serum Replacement (KOSR), 1x Non-essential Amino Acids (NEAA, Gibco), 1x Glutamax (Gibco)). After 24h in simple transition medium (day 0), medium was exchanged for BAP medium (DMEM/F12, 20% KOSR, 1x NEAA, 1x GlutaMax, 10ng/mL bone morphogenic protein 4(BMP4), 1µM of the TGFβ inhibitor (A83-01), and 0.1µM of an FGF receptor inhibitor (PD173074). BAP medium was exchanged daily. After 72h in BAP medium (day 3), cells were ∼90-100% confluent and were passaged following dissociation with TrypLE Express (manufacturer) for 8 minutes. TypLE was quenched with BAP medium, cells were collected and pelleted for 3 minutes at 150 rcf. Supernatant was removed and cells were resuspended in BAP medium and replated in Matrigel coated wells at ∼100,000 cells/9.5cm^2^. Medium was exchanged daily with fresh BAP medium before fixation/collection at day 8. Cells were then fixed with 4% paraformaldehyde from 20 minutes or single- cell digested with TrypLE for 5 min, quenched with BAP medium, and pelleted via centrifugation for downstream analysis.

#### RNA Isolation

Total RNA was extracted from frozen cell pellets with the New England Biolabs Monarch on-column RNA extraction kit according to manufacturer protocol and was treated with DNAse1 on-column to remove genomic DNA. RNA quality was measured on a Nanodrop (ThermoFisher Scientific) by spectrophotometry and only samples with a 260/230 ratio >1.7 and 260/280 ratio of >1.7 were used for downstream experiments. RNA integrity was measured on a Tape Station (Agilent) and was evaluated for clear 18S and 28S rRNA bands (RIN>9) for RNA-sequencing experiments.

#### RNA-sequencing and Analysis

One individual replicate for each of 6 biological replicates (iPS cell lines) per ploidy status (T21 or D21) were used for RNA-seq experiments. Additionally, we used 3 individual replicates of the isogenic iLD11 T21 and D21 cell lines for additional RNA-seq experiments. To generate libraries for bulk RNA-seq experiments, 200ng of high-quality RNA was prepared with the Universal Plus mRNA-Seq library preparation kit with NuQuant (Tecan). Paired-end sequencing reads of 150bp were generated on a NovaSeqX (Illumina) sequencer at a target depth of 80 million paired-end reads per sample. Raw sequencing reads were de-multiplexed using bcl2fastq.Fastq files were processed with the rnaseq Nextflow computational pipeline [version number 3.17.0, (52)] on the University of Colorado supercomputer cluster. Raw counts were analyzed to produce differentially expressed gene tables with DESeq2 in RStudio (version 2026.01.1+403) with submitted batch, aneuploidy status, and age/sex-matched pairs as variables. Lowly expressed genes (<10 counts in at least 3 cells) were filtered out prior to DESeq2.

For Gene Set Enrichment Analysis de novo trophoblast gene sets, the top 1000 differentially expressed genes by Log2FC after DESeq2 were used from publicly available RNA-seq data (25). For enrichment of D21 and T21 differentially expressed genes in publicly available single-cell data from placental tissue, we entered T21/D21 gene lists into the Placenta Cell Enrich (53) and Reproductive Cell Atlas (54) tools to generate figures.

#### Quantitative RT-PCR

500ng of RNA was used to make cDNA with the New England Biolabs LunaScript RT Supermix. Quantitative RT-PCR was performed with New England Biolabs Luna qPCR mastermix in triplicate on an Agilent Biosystems 7500 qPCR machine. Relative transcript expression was quantified with the 2^ΔΔCt^ method. Primers are listed in Table 1.

#### Immunofluorescence and Imaging

BAP-differentiated cells and pre-implantation blastoids were fixed with 4% paraformaldehyde for 20 min. Cells were then washed with 0.1% Tween and 0.1% polyvinyl-pyrrolidone in phosphate-buffered saline (PBST) three times for 5 min. Cells were permeabilized with 0.3% Triton-X in PBS for 30 min, washed 3 times in PBST for 5 min, and blocked with 0.1% Tween, 1% bovine serum albumin (BSA), 0.1M glycine, and 10% horse serum for 2h. Primary antibodies (1:200) were left on overnight. Cells were washed three times with PBST and stained with secondary antibodies (1:200) for 1h.

#### Fusion Index Quantification

Cells quantified for fusion indices were stained for antibodies against a cell border marker (ZO-1) or with rhodamine phalloidin to stain F-actin and DAPI for nuclei. Images were taken at 40X magnification on a fluorescence microscope. Fusion indices was manually quantified as ((# nuclei within a multinucleated syncytia- # of syncytia)/ total nuclei) per field of view. All fusion index images were blinded before quantification. Fusion index experiments were performed with at least 10 frames for each of 3 individual replicates per cell line. Estrogen supplementation experiments were performed with 40 frames across 4 individual replicates (T21/D21) or 60 frames across 3 individual replicates to overcome the variance from the treatment groups.

## Acknowledgments

We would like to thank the Linda Crnic Institute for Down Syndrome and the Global Down Syndrome Foundation for their support in deriving and providing iPS cell lines and funding our RNA-sequencing experiments. Blastoid generation was funded by the Anna and John J. Sie Foundation through a Postdoctoral Sie Fellowship and with startup funds from the University of Colorado. We are grateful for funds for all other experiments from NIH NRSA F32 award number 1F32HD116568-01 and NIH R21 award number 1R21HD118444-01. The iPS cell lines from the Human Trisome Project are maintained and distributed with funding from the NIH INCLUDE Project through a grant from the National Institute on Aging (grnat U24AG092191). We would also like to acknowledge the Light Microscopy Core Facility, Porter Biosciences B047, B049, B051 and B059 at the University of Colorado Boulder (RRID:SCR_018993) for help and advice with microscopy.

## Author contributions

D.M.L. performed experiments and wrote the manuscript with input from all authors. I.T.P. developed the NRIP1 overexpression iPS cell line. B.B., K.L.W., L.S., M.N., L.O., B.D.T. and C.H. assisted in data collection or analysis. B.N. maintains donor-derived iPS cells. R.D., M.A, J.R., J.M.E., M.D.G., and D.J.S. advised the study and assisted with manuscript preparation. J. B. and D.M.L. conceived the study and designed experiments.

## Figure legends

**Figure S1.**
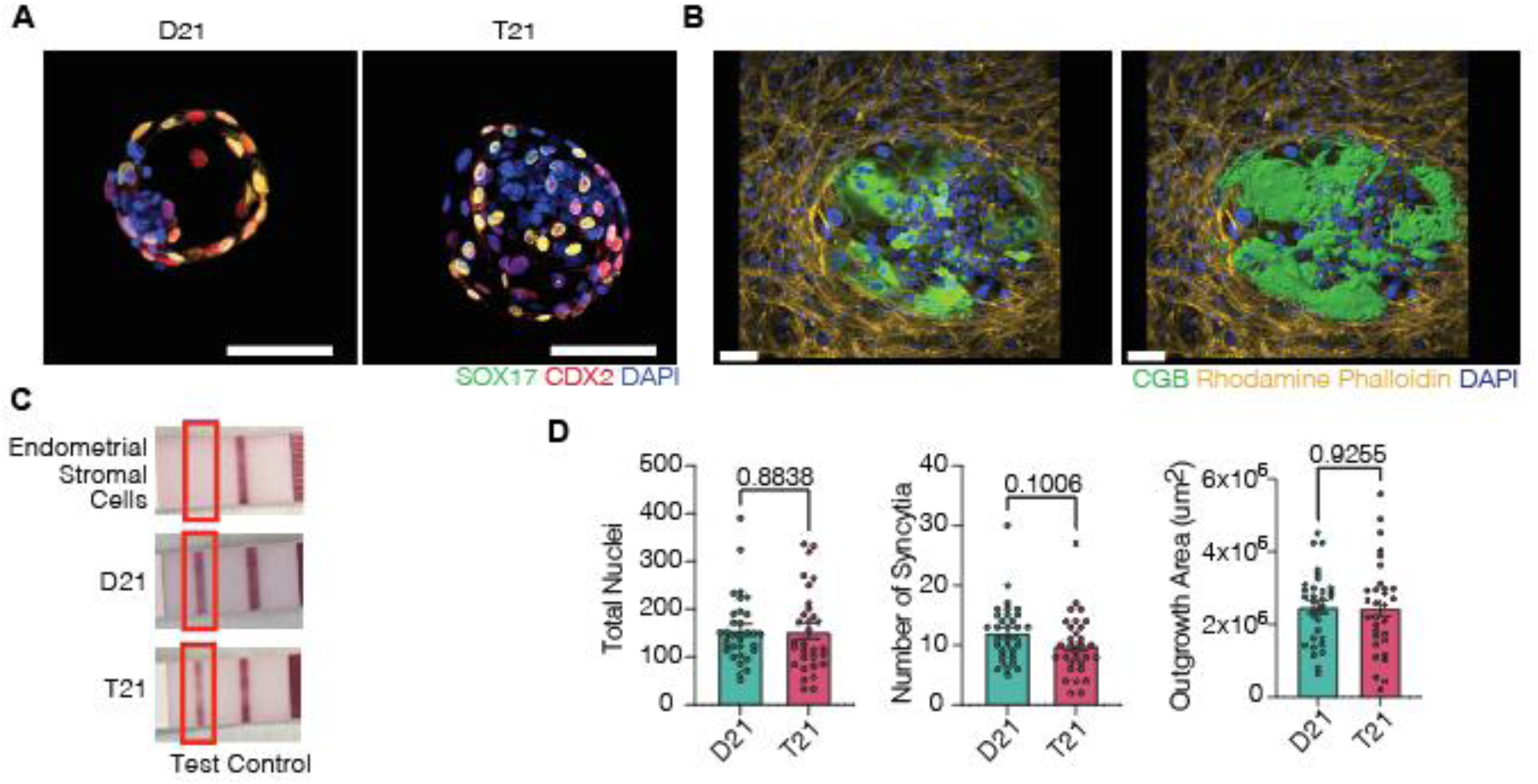
A) Representative immunofluorescence images of T21/D21 blastoids stained for antibodies against SOX17 (green), CDX2 (red), and DAPI (blue). Scale bar=100µm. B) Representative immunofluorescence images of blastoids implanted for 72h stained for antibodies against human chorionic gonadotropin subunit beta (CGB, green) rhodamine phalloidin=yellow, DAPI=blue. Scale bar=100µm. Right panel shows CGB+ volume by surface tool in Imaris. C) Pregnancy test strips for secreted human chorionic gonadotropin from spent medium for immortalized endometrial stromal cells, or D21/T21 blastoids following 72h of implantation onto immortalized endometrial stromal cells. D) Total nuclei, number of syncytia, and outgrowth area, for blastoids implanted after 72h. Data are represented as mean ± SEM. P-values are shown following student **t** test. n=9 blastoids for each of 4 individual replicates.

**Figure S2.**
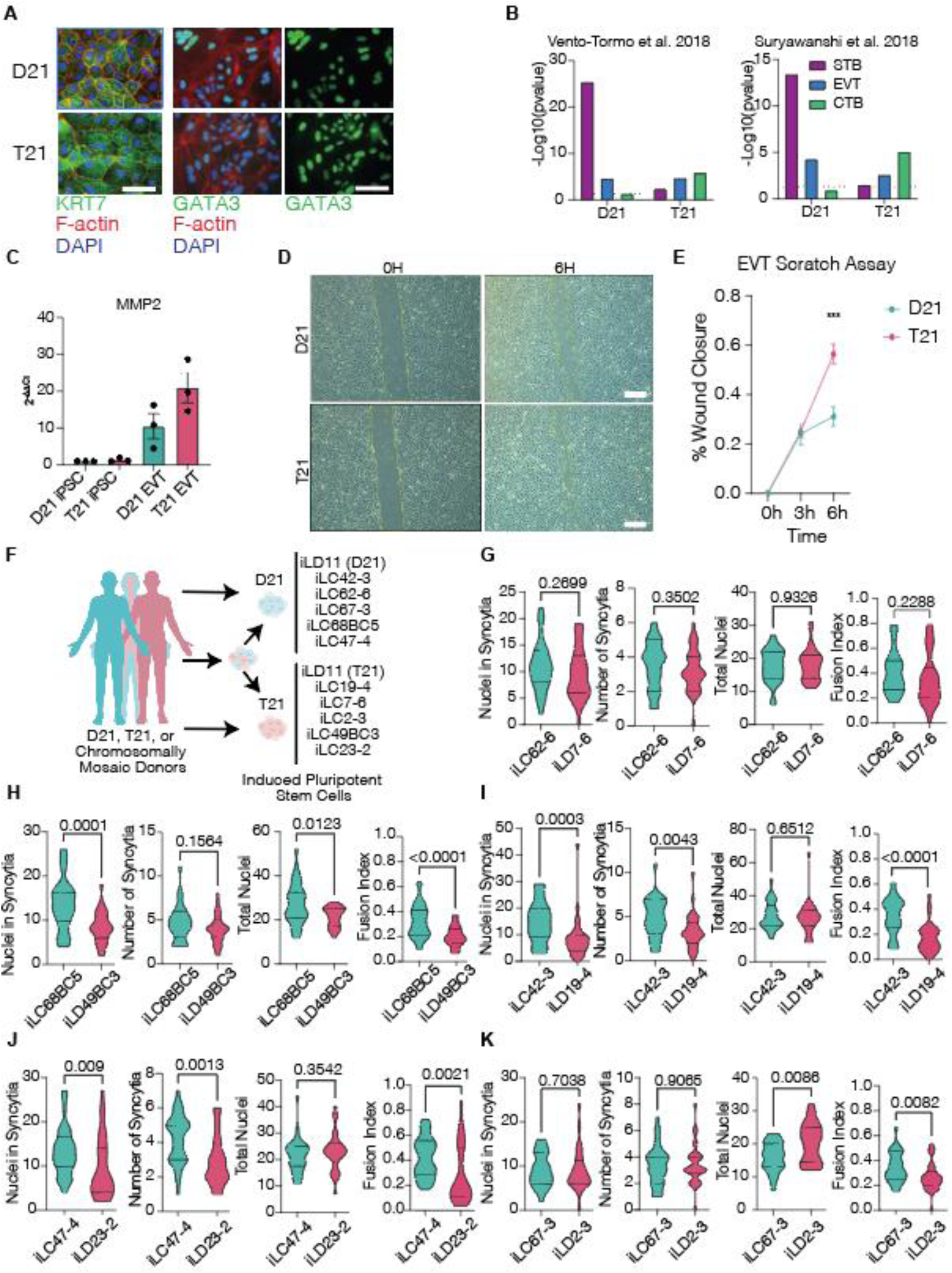
A) Representative immunofluorescence images of T21/D21 trophoblast-like cells stained for antibodies against trophoblast markers KRT7 (green, left) or GATA3 (right), rhodamine phalloidin (red, left only), and DAPI (blue, left only). Scale bar=100µm. B) Enrichment of syncytiotrophoblast, extra-villous trophoblast, and villous cytotrophoblast gene sets in D21 or T21 isogenic samples(26,27). C) RT-qPCR for EVT gene MMP2 in EVT-like cells compared to undifferentiated iPS cells. D) Representative brightfield images of a scratch wound assay in isogenic T21/D21 EVT-like cells after 0h and 6h. Scale bar= 100µm. E) Quantification of % wound closure in T21/D21 isogenic EVT-like cells. ***p<0.001 after student T. test. Data are represented as mean ± SEM. n=3 technical replicates (scratches) for each of 3 individual replicates. F) Schematic for T21, D21, or chromosomally mosaic donor-derived iPS cells. G) Nuclei in syncytia, number of syncytia, total nuclei, and fusion index for age- and sex-matched pair iLC62-6 and iLD 7-6, H) iLC68BC5 and iLD49BC3, I) iLC42-3 and iLD19-4, J) iLC473 and iLD23- 2, and K) iLC67-3 and iLD2-3. P-values are shown following student T. test. n= 30 fields of view across 3 individual replicates for each cell line.

**Figure S3.**
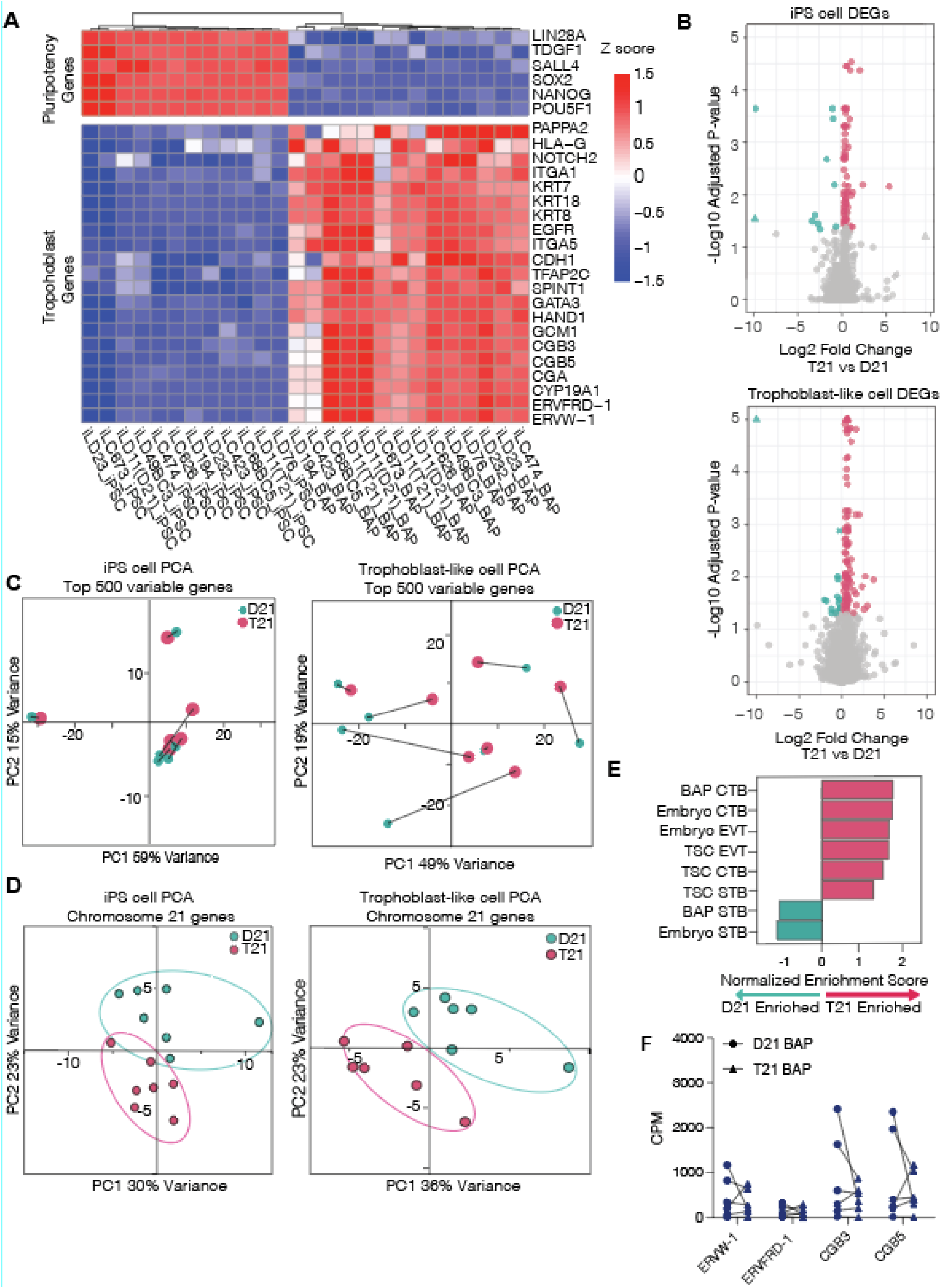
A) Heatmap with Z-score scaled expression of pluripotency factors and trophoblast markers in all D21/T21 undifferentiated induced pluripotent stem cells (iPSC) and trophoblast-like cells (BAP). B) Volcano plot highlighting significantly (adjusted p<0.05) differentially expressed genes with color between T21 (>0 Log2 fold change) and D21 (<0 Log2 fold change) in undifferentiated iPSC (left) or trophoblast-like cells (right). C) PCA for all age- and sex- matched pairs in undifferentiated iPSC (left) or trophoblast-like cells (right) for top 500 variable genes. D) PCA for all age- and sex- matched pairs for genes on chromosome 21 in undifferentiated iPSC (left) or trophoblast-like cells (right). E) GSEA for CTB, STB, and EVT gene sets from peri-implantation embryos(West et al., 2019)(West et al., 2019) BMP4-treated trophoblast-like cells (BAP), and trophoblast stem cells (TSC) (Logsdon et al., 2024b)(Logsdon et al., 2024b) with normalized enrichment score in all 12 T21/D21 trophoblast-like samples. F) Transcript abundance in counts per million mapped read (CPM) for STB genes *ERVW-1*, *ERVFRD-1*, and *CGB3/5* in age-and sex-matched pairs. Pairs are connected by a line. Pairs are connected by a line. *p<0.05, **p<0.01 after student T. test. n= 1 individual replicate each of 6 biological replicates for each ploidy (T21 and D21).

**Figure S4.**
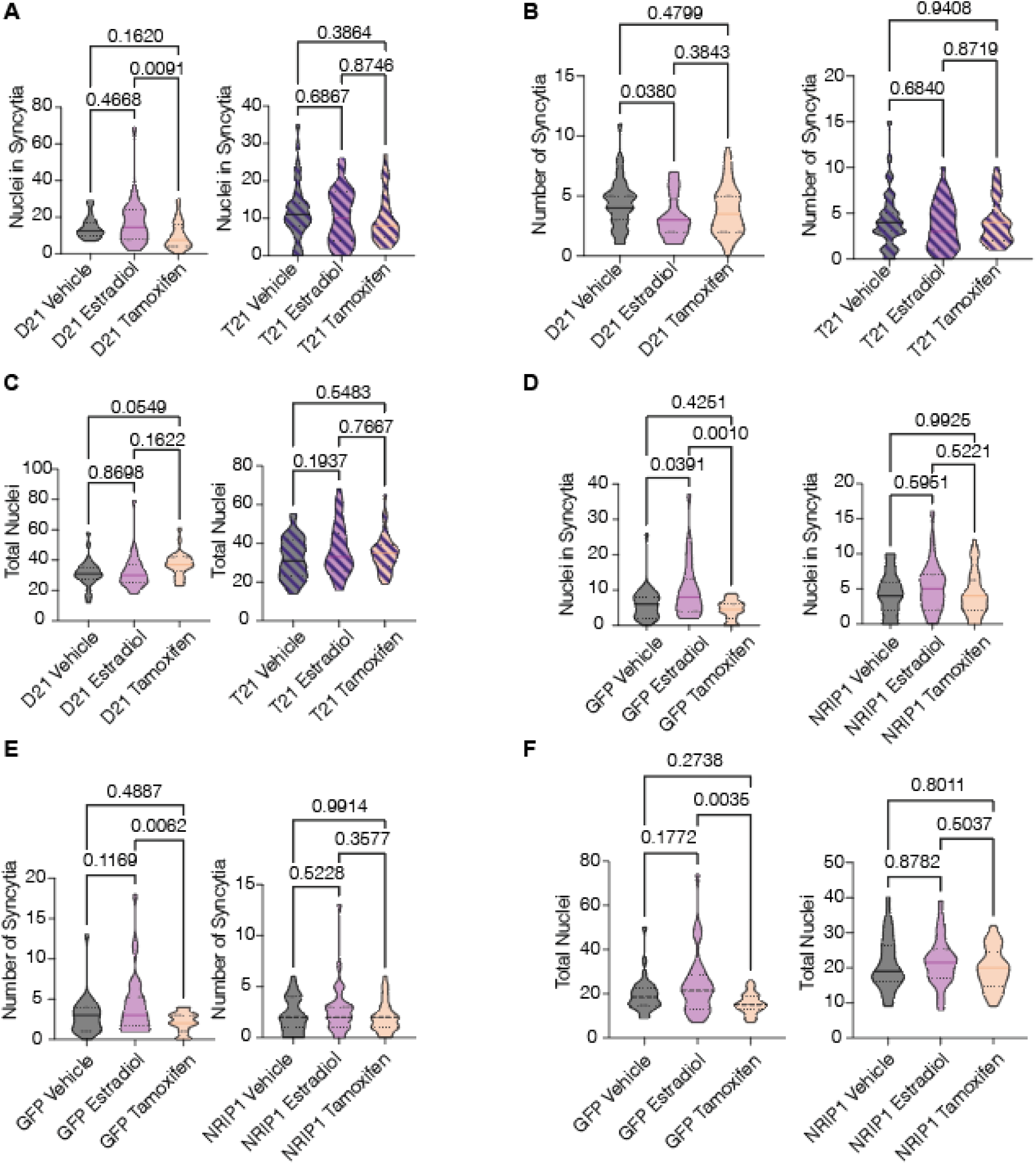
A) Nuclei in syncytia, C) number of syncytia, and D) total nuclei in T21/D21 trophoblast-like cells following treatment in vehicle, 1 µM 17β-estradiol, or 1µM Tamoxifen. E) Nuclei in syncytia, F) number of syncytia, and G) total nuclei in GFP/NRIP1 overexpression trophoblast-like cells following treatment in vehicle, 1 µM 17β-estradiol, or 1µM Tamoxifen. P-values are noted following a one-way ANOVA with Tukey’s test for multiple comparisons.

